# Theseus: Fast and Optimal Affine-Gap Sequence-to-Graph Alignment

**DOI:** 10.64898/2026.02.12.705572

**Authors:** Albert Jiménez-Blanco, Lorién López-Villellas, Juan Carlos Moure, Miquel Moreto, Santiago Marco-Sola

**Affiliations:** Department of Computer Science, Universitat Politècnica de Catalunya, Barcelona, Spain; Computer Sciences Department, Barcelona Supercomputing Center, Barcelona, Spain; Department of Computer Architecture and Operative Systems, Universitat Autònoma de Barcelona, Barcelona, Spain; Department of Computer Science and Systems Engineering, University of Zaragoza, Spain; Department of Computer Architecture, Universitat Politècnica de Catalunya, Barcelona, Spain

## Abstract

**Motivation:** Sequence-to-graph alignment is a central problem in bioinformatics, with applications in multiple sequence alignment (MSA) and pangenome analysis, among others. However, current algorithms for optimal affine-gap alignment impose high memory and computational requirements, limiting their scalability to aligning long sequences to complex graphs. Practical solutions partially address this problem using heuristic strategies that ultimately trade off optimality for speed.

**Results:** This work presents Theseus, a novel, fast, and optimal affine-gap sequence-to-graph alignment algorithm. Theseus leverages similarities between genomic sequences to accelerate the alignment computation and reduces the overall memory requirements without compromising optimality. To that end, Theseus exploits the diagonal transition property to process only a subset of the dynamic programming cells, combined with a sparse-data strategy that enables efficient sequence-to-graph alignment. Moreover, our algorithm supports optimal affine-gap alignment on arbitrary directed graphs, including those with cycles. We evaluate Theseus on two key problems: multiple sequence alignment (MSA) and pangenome read mapping. For MSA, we compare it against the state-of-the-art methods SPOA, abPOA, and POASTA. Theseus is 2.0*×* to 232.2*×* faster than the other two optimal aligners, SPOA and POASTA. Compared with abPOA, a heuristic aligner, Theseus is 3.3*×* faster on average, while ensuring optimality. For pangenome read mapping, we benchmark Theseus against the alignment stage of the popular mapping tool vg map, along with the alignment kernels of SPOA, abPOA, and POASTA. Theseus outperforms the other methods, showing a 1.9*×* to 16.9*×* speed improvement on short reads.

**Availability:** Theseus code and documentation are publicly available at https://github.com/albertjimenezbl/theseus-lib.

## 1 Introduction

Sequence-to-graph alignment is a central problem in modern bioinformatics (Eizenga *et al*., 2020) driven by the growing use of sequence graphs to compactly model genetic diversity (Miga and Wang, 2021; com, 2018) and reduce reference bias (Garrison *et al*., 2018). Consequently, sequence-to-graph alignment has applications across multiple fields, including comparative genomics (Tettelin *et al*., 2008), evolutionary biology (Zhang *et al*., 2019), and pangenomics (Liao *et al*., 2023), among others (Compeau *et al*., 2011; Miller *et al*., 2010; Marco-Sola *et al*., 2012; Marco-Sola and Ribeca, 2015), enabling fundamental tasks such as Multiple Sequence Alignment (MSA) (Lee *et al*., 2002) and read mapping to pangenome graphs (Rautiainen and Marschall, 2020).

Multiple Sequence Alignment (MSA) is a fundamental analysis in bioinformatics and genomics (Gotoh, 1999), providing the basis for studying conservation, variation, and evolutionary relationships across sequences. To that end, Partial Order Alignment (POA) (Lee *et al*., 2002) is a widely used method that progressively builds an alignment of a set of *N* sequences. In its original formulation, POA provides an efficient and informative solution to the MSA problem (Lee *et al*., 2002), progressively building a partial order graph that represents the incremental multiple alignment. This is done by iteratively aligning each new sequence *S*_*i*_ to the current sequence-graph and then updating the graph to incorporate each newly computed alignment.

Similarly, pangenome read mapping has emerged as a fundamental problem in bioinformatics, crucial for analyzing genetic variation across populations by replacing traditional linear reference with pangenome graphs. Initiatives such as the Human Pangenome Reference Consortium (Liao *et al*., 2023) and the 1000 Genomes Project (Consortium *et al*., 2015) reflect the growing interest in efficiently modeling genetic diversity and reducing reference bias in the analysis of highly variable regions, such as the Major Histocompatibility Complex (MHC) (Chin *et al*., 2020). Similar to traditional read-mapping tools, most currently used pangenome read mappers (Garrison *et al*., 2018; Rautiainen and Marschall, 2020) implement the seed-and-extend paradigm. This paradigm consists of two phases: (1) A seeding strategy is used to find candidate mapping locations, and (2) These candidates are extended using an alignment algorithm to determine the better-scoring locations in the reference pangenome graph. Current pangenome read mappers usually work with directed acyclic graphs (DAG) (Garrison *et al*., 2018) or admit small cycles (Rautiainen and Marschall, 2020), but do not directly support large cycles. VG implicitly represents such cycles by unfolding the cyclic components in the reference pangenome. However, this unfolding process can substantially increase the size of the unfolded graph with respect to its cyclic counterpart (Kavya *et al*., 2019).

However, the large scale of sequence-graphs used for MSA and pangenome analyses, combined with the ever-growing volume of genomic data (Goodwin *et al*., 2016), places significant stress on traditional sequence-to-graph alignment algorithms (com, 2018; López-Villellas *et al*., 2024, 2023). Even for the simplified version of DAGs, the algorithmic cost of traditional algorithms grows proportionally to the product of the number of bases in the reference graph and the length of the aligned sequence (Jain *et al*., 2020). The problem becomes harder when considering cyclic graphs. As a result, traditional methods require high memory and runtime and scale poorly when dealing with large numbers of sequences or complex, large pangenome graphs (Darby *et al*., 2020).

Classical solutions rely on extending the dynamic programming (DP) formulation of sequence-to-sequence alignment (Needleman and Wunsch, 1970) to sequence-graphs. Navarro introduced the first such extension, proposing a DP algorithm that computes the optimal alignment of a sequence to an arbitrary graph for approximate hypertext matching (Navarro, 2000). This foundational algorithm was later extended to support arbitrary costs (Jain *et al*., 2020). However, its time and memory demands make it impractical for large-scale genomic data, which has motivated the proposal of multiple constant-factor optimizations, such as SIMD vectorization and bit-parallel techniques (Myers, 1999; Doblas *et al*., 2023, 2025b). Notably, SPOA (Vaser *et al*., 2017) is a widely used state-of-the-art POA library for computing MSA that speeds up the traditional DP computation through SIMD vectorization. Similarly, GraphAligner (Rautiainen *et al*., 2019; Rautiainen and Marschall, 2020) is a widely used pangenome read-mapping tool that speeds up edit-distance computation through bit-parallel techniques.

In contrast to optimal approaches, many practical implementations employ aggressive heuristics to accelerate computation and reduce the memory usage at the expense of accuracy, often missing the optimal alignment solution. Examples of such heuristic strategies include the banded alignment mode of GraphAligner (Rautiainen and Marschall, 2020), the seed-chaining approach of minigraph (Li *et al*., 2020), and the adaptive banding in abPOA (Gao *et al*., 2021).

More recently, new output-sensitive methods (Doblas *et al*., 2025a) have been proposed whose runtime scales with the difficulty of the alignment (i.e., alignment score), adjusting to the workload complexity. Notably, the Wavefront Alignment algorithm (WFA) (Marco-Sola *et al*., 2021, 2023; Aguado-Puig *et al*., 2023; Haghi *et al*., 2023; Alonso-Marín *et al*., 2024) introduced an output-sensitive formulation that achieves optimal sequence-to-sequence alignment, but it is limited to linear references. Other methods, inspired by classical informed searches (Hart *et al*., 1968), implement *A*^∗^-based strategies to reduce the number of DP cells explored in graph alignment. POASTA (van Dijk *et al*., 2025) and Astarix (Ivanov *et al*., 2020) follow this strategy, reducing DP cell computations at the expense of overheads for maintaining a priority queue, computing the heuristic function, and requiring additional memory.

In this work, we present Theseus, a fast and memory-efficient output-sensitive algorithm for optimal affine-gap sequence-to-graph alignment. Inspired by the WFA algorithm for sequence-to-sequence alignment (Marco-Sola *et al*., 2021, 2023), Theseus exploits the similarity between sequences to accelerate computations and reduce memory footprint. Unlike other sequence-to-graph alignment algorithms tailored for specific use cases, Theseus is flexible enough to be applied to many sequence-to-graph alignment applications. Moreover, Theseus works on directed graphs and supports cycles, efficiently addressing the sparse nature of graph exploration.

We evaluate Theseus on two fundamental and critical applications: MSA computation and pangenome read mapping. First, for MSA computation, we compare Theseus against the state-of-the-art POA methods SPOA and POASTA (exact) and abPOA (heuristic). Our results show that Theseus is 2.0*×* to 232.2*×* faster than the other exact aligners. Moreover, Theseus is 3.3*×* faster on average than abPOA, while ensuring optimality. Second, for read mapping, we compare Theseus against the sequence-to-graph alignment kernels in vg map, SPOA, abPOA, and POASTA. In this experiment, Theseus outperforms the other methods, showing a 1.9*×* to 16.9*×* speed improvement on short reads. In summary, this work makes the following contributions.

- We present Theseus, a novel output-sensitive algorithm for fast and memory-efficient computation of optimal affine-gap sequence-to-graph alignment that supports reference graphs with cycles.
- We propose a diagonal invalidation strategy that compactly represents and updates the set of invalid alignment diagonals, eliminating redundant computations during sequence-to-graph alignment.
- We introduce sparse wavefronts, an efficient data structure to store intermediate wavefronts over sequence-graphs. We also propose a sequence-graph unified coordinate system to efficiently operate over multiple sparse wavefronts across the vertices of a sequence-graph.
- We introduce a memory-efficient backtrace algorithm to recover the optimal alignment path from sparse wavefronts over sequence-graphs.
- We provide a high-performance, open-source implementation of Theseus, distributed as both a library and two stand-alone minimal tools, publicly available to the community.

## 2 Background

A sequence-graph is a tuple *G* = (*V, E, σ*), where *V* denotes a set of vertices, *E* ⊆ *V × V* its set of directed edges, and *σ* : *V* → Σ^∗^ assigns a sequence of symbols over alphabet Σ to every vertex. For simplicity, we denote the length of the sequence associated with *v* as |*v*|. Given vertices *s, t* ∈ *V*, a path *P* from *s* to *t* is defined as the ordered sequence of vertices (*s, v*_1_, …, *v*_*k*_, *t*) connected by edges in the sequence-graph *G*. Any path *P* = (*s, v*_1_, …, *v*_*k*_, *t*) has an associated sequence-path *S*_*P*_ = *σ*(*s*) · *σ*(*v*_1_) … *σ*(*v*_*k*_) · *σ*(*t*) obtained by concatenating the sequences of its vertices. Let *Q* be a sequence of length *m*, which we name *query*, and let *G* be a sequence-graph, which we name *graph-reference*. Then, the optimal sequence-to-graph alignment is the sequence of four basic operations (i.e., match, mismatch, insertion, and deletion) that transform *Q* into a sequence *S*_*P*_, spelled by a path in *G* originating at some vertex *s*, while minimizing a given score function.

Most practical applications employ score function models: gap-linear and gap-affine. The first is the simple gap-linear score function (also known as weighted edit distance), which assigns fixed penalty values (*a, x, e*) for matches, mismatches, and gap extensions (applied to both insertions and deletions), respectively. The second model is the affine-gap score function (Gotoh, 1982), defined as a tuple (*a, x, o, e*) of four values, where *a* is the cost of a match, *x* is the cost of a mismatch, *o* is the cost of opening (starting) a gap and *e* is the cost of extending it. This formulation models the fact that starting a new gap is more expensive than extending an existing one. For example, two insertions interrupted by another operation require paying two gap-opening and two extension penalties 2*o* + 2*e*, while a single uninterrupted two-base gap costs only *o* + 2*e*. Using a gap-affine function allows treating long gaps as a single event rather than multiple independent edits, yielding more accurate biological interpretations.

The optimal sequence-to-graph alignment can be computed using the recurrence equations in Eq. 1. *I*_*v,i,j*_ and *D*_*v,i,j*_ represent the optimal score to align the prefix *Q*[1, *i*] against any valid path that reaches offset *j* of vertex *v* and ends in an insertion or a deletion, respectively. The third value, *M*_*v,i,j*_, represents the optimal alignment score of reaching position (*v, i, j*), regardless of the last edit operation. The initial conditions (*j* = 0) at each vertex *v* depend on the last scores computed at every vertex *u* (*j* = |*u*|) with an incoming edge to *v*; i.e, (*u, v*) ∈ *E*.

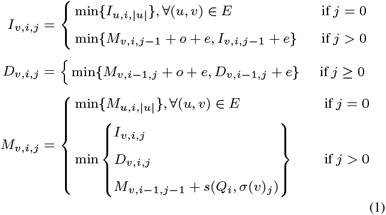

In practice, solutions to Eq. 1 are computed using DP, as the problem exhibits optimal substructure where every partial solution of an optimal alignment is optimal. Common implementations under the affine-gap model allocate three DP matrices per vertex *v* (*M*_*v*_, *I*_*v*_, and *D*_*v*_) of dimension *O*(*m*|*v*|). This results in *O*(*m*(*L* + |*E*|)) time and space complexity (Navarro, 2000), with *L* = ∑_*v*∈*V*_ |*v*| the total length of the sequences in the graph.

### 2.1 Wavefront Alignment Algorithm

For the sequence-to-sequence alignment problem, the Wavefront Alignment Algorithm (WFA) (Marco-Sola *et al*., 2021) is an output-sensitive algorithm that leverages sequence similarity to accelerate the alignment computation. Unlike traditional DP-based algorithms, which require computing the full DP matrix, WFA evaluates DP cells in order of increasing alignment score until the optimal solution is found. Specifically, WFA computes partial alignments for increasing values of score, from *s* = 0, until the optimal alignment score *s*^∗^.

The WFA algorithm relies on two key concepts: diagonal-encoding and furthest-reaching cells. First, to compute the optimal alignment between two sequences *Q* and *T*, any DP cell (*i, j*) in the matrix can be uniquely identified by its *diagonal k* = *j* −*i* and its offset *off* = *i*, which specifies the position of the cell along that diagonal. This defines a coordinate transformation from (*i, j*) to (*k, off*), where the original indices can be recovered as (*i, j*) = (*off, k* + *off*). Second, for a given score *s* and diagonal *k*, the *furthest-reaching cell* ℱ_*s,k*_ represents the cell with score *s* on diagonal *k* with the most advanced offset *off* along the diagonal. The vector of all furthest-reaching cells across all diagonals for a given score *s* is called the *wavefront* at score *s*. Focusing only on the furthest-reaching cells for each score and diagonal, WFA avoids computing the full DP matrix, resulting in significant efficiency gains for similar sequences.

Under the affine-gap score model, WFA computes the optimal sequence-to-sequence alignment using the recurrence relations in Eq. 2. To that end, WFA maintains three wavefront data structures, 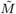, *Ĩ*, and 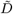, which record all the wavefronts for each score *s*, corresponding to the match-mismatch state, the insertion state, and the deletion state of the DP formulation, respectively. The algorithm proceeds iteratively by score, alternating between two phases: *next* and *extend*. In the *next* phase, for each score *s*, the algorithm computes the initial offsets of 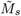, *Ĩ*_*s*_, and 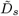 for all reachable diagonals by applying Eq. 2 using previously computed wavefronts. Then, during the *extend* phase, each initial furthest-reaching cell in the 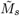 wavefront is advanced following consecutive matches along the diagonal (i.e., computing the Longest Common Prefix, LCP, between the corresponding suffixes of the query *Q* and the text *T*). The algorithm terminates when it first meets the alignment’s end condition, immediately identifying the optimal score *s*^∗^.

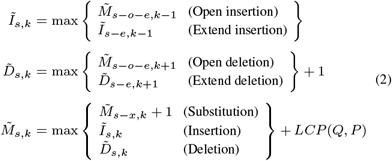

As a result, WFA computes *s* wavefronts of increasing size and evaluates an *LCP* () operation in each of them. Each diagonal offset can only advance up to the length of the sequences. Thus, the time complexity of WFA is *O*(*ns*) when the sequences are similar (*s* ≪ *n*).

## 3 Methods

In this section, we describe our method, Theseus, to compute fast and optimal affine-gap sequence-to-graph alignment. We begin by generalizing the notion of furthest-reaching cells to sequence-graphs and deriving the corresponding recurrence relations (Section 3.1). Then, we present the Theseus algorithm, highlighting the particular challenges that arise when aligning against a sequence-graph reference. (Section 3.2). Next, we introduce a diagonal invalidation strategy that avoids redundant work by tracking explicitly and implicitly explored diagonals (Section 3.3). Then, we describe a unified coordinate system that enables the efficient combination and processing of sparse wavefronts from different vertices (Section 3.4). Finally, we present a memory-efficient backtrace method to reconstructs the optimal alignment path from the sparse wavefront representation (Section 3.5).

### 3.1 Furthest-Reaching Cells in Sequence-Graphs

Theseus generalizes the core concepts of the WFA algorithm to the broader problem of sequence-to-graph alignment, progressively exploring the solution space by computing partial alignments with increasing score. For that, Theseus computes successive wavefronts of furthest-reaching cells using Eq. 3.

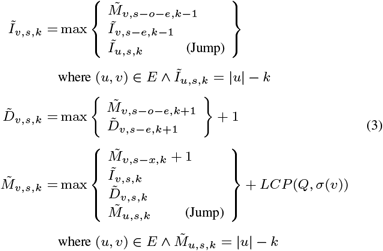

We propose to work with three vertex-local wavefront structures 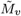, *Ĩ*_*v*_, and 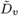 for each vertex *v* ∈ *V*, instead of using three global structures for all vertices. This way, for each score *s*, vertex *v*, and diagonal *k*, Theseus computes the furthest-reaching cell as a maximum over the possible previous states reaching the position (*v, s, k*) in 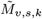, *Ĩ*_*v,s,k*_, and 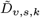.

However, note that recurrences on Eq. 3 are not vertex-local, as their values may depend on wavefront values from adjacent vertices in the sequence-graph. This requires redefining the recurrence equations to account for jumps across vertices, ensuring that the furthest-reaching cells at score *s* include contributions from incoming wavefronts. Note that a diagonal *k* in vertex *v* can be reached in two different ways: either from a diagonal in the same vertex *v* or from a diagonal of an incoming vertex *u* s.t. (*u, v*) ∈ *E*. Therefore, *Ĩ*_*v,s,k*_ and 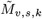 must also account for the offsets propagated from each incoming vertex *u* where the diagonal *k* in *u* has an offset value of |*u*| − *k*; that is, *Ĩ*_*u,s,k*_ = |*u*| − *k* and 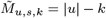, respectively.

### 3.2 Theseus Algorithm

Theseus’ algorithm explores the solution space in increasing order of score *s* using Eq. 3. It starts at an initial vertex *v*_0_ with score *s* = 0 and continues until it reaches a path in *G* that yields a complete alignment of *Q*. Since this is the first complete alignment of *Q* found during its score-ascending search, it is necessarily optimal and achieves the minimal alignment score *s*^∗^. Algorithm 1 presents the overall structure of the Theseus algorithm.

At every iteration, the algorithm maintains a set of active vertices *V*_*act*_, a set of active diagonals *K*_*act*_, and a set of invalidated diagonals *K*_*inv*_. We say that a vertex *v* is active if it has at least one active diagonal, where an active diagonal is defined as a diagonal in *v* that can be reached with score *s* and has not yet been invalidated. Thus, the active diagonals represent the positions within active vertices where the alignment can still progress. Limiting computation to this active set allows the algorithm to focus its effort on the regions of the graph that directly influence the alignment, improving both time and memory efficiency, as can be seen in Figure 1. Each time a diagonal *k* reaches the end of a vertex, the diagonal is added to the set *K*_*inv*_(*v*) of invalid diagonals in vertex *v*. An efficient implementation of this method, accounting for the implicit invalidation of diagonals due to gap propagation, is explained in Section 3.3.

**Fig. 1:**
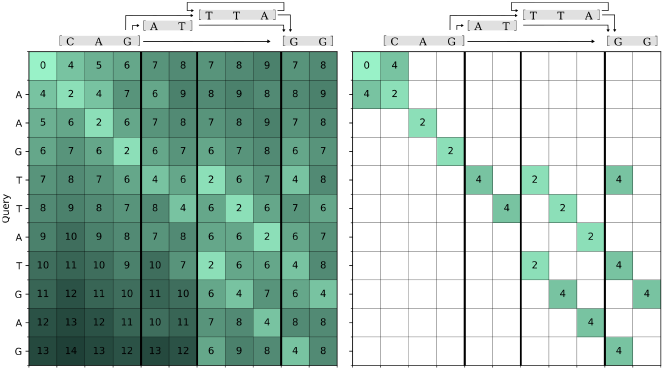
Comparison of the cells explored by traditional DP methods and the Theseus algorithm on a graph containing a cycle.

The alignment process begins by setting initial conditions. In the case of a global alignment starting at the vertex *v*_0_, it suffices to set 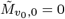 (score *s* = 0, first wavefront). Initially, the set of active vertices is set to *V*_*act*_ = *{v*_0_*}*, the set of active diagonals is 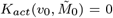, and the set of invalid diagonals *K*_*inv*_ is empty. The set of active vertices and diagonals is updated in each iteration due to jumps and the opening of new diagonals resulting from insertions and deletions, respectively. Then, for each score *s*, starting at *s* = 0, Theseus performs the two core operations, *next* and *extend*, until the end condition is met at score *s*^∗^.

#### Algorithm 1

Theseus sequence-to-graph alignment algorithm

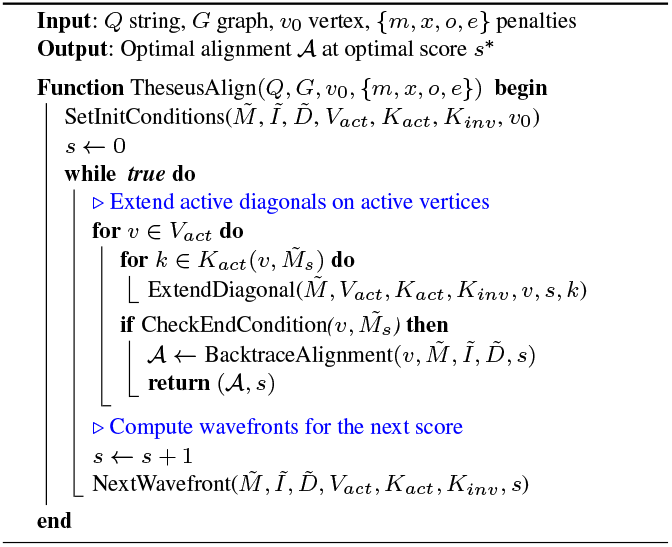

On each iteration of Algorithm 1, Theseus begins by extending each active diagonal (Algorithm 2). To that end, every active offset 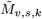 is greedily extended within its vertex *v* by computing the LCP between the suffixes 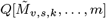 and 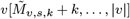. If the extension reaches the end of the vertex, that is, 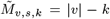, the diagonal *k* is invalidated in vertex *v*, and the algorithm proceeds to process every outgoing vertex *u* from *v*. Given a vertex *v*, we name *v*_*out*_ = *{u* ∈ *V* | (*v, u*) ∈ *E}* the set of its outgoing vertices. For every *u* ∈ *v*_*out*_, *u* and *k* are marked as active (if they weren’t yet) and the value at 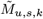 diagonal *k* is updated. If the diagonal *k* at vertex *u* is inactive, the extension continues within *u*. The algorithm repeats this procedure until it finds a mismatch. Most importantly, the algorithm makes no assumptions about the graph topology and correctly handles graphs containing cycles.

#### Algorithm 2

ExtendDiagonal to advance diagonals

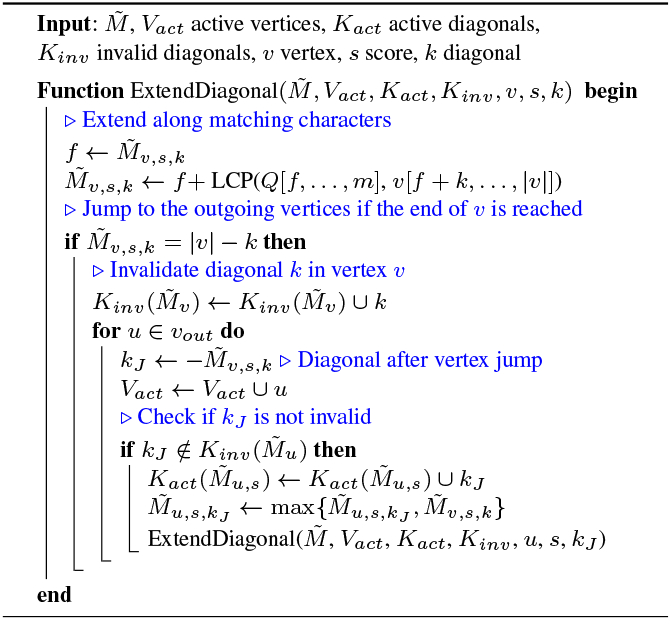

After extending all active diagonals, Theseus evaluates the end condition associated with the problem at hand. In the case of global alignment with starting at vertex *v*_0_, for instance, the algorithm satisfies the end condition the first time the query *Q* is completely processed. When Theseus meets this condition for the first time, it traces back the path of an optimal alignment and returns it to the user, along with the optimal score *s* = *s*^∗^.

If the end condition is not met, Theseus computes wavefronts with the next score *s* = *s* + 1. For each of the active vertices *v*, the *Ĩ*_*v,s*_, 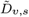, and 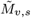 wavefronts are computed based on the values at the corresponding wavefronts with a lower score, according to Eq. 3. In particular, each time that Theseus reaches the end of a vertex *v* with an insertion, it invalidates the corresponding diagonal and continues the alignment in its outgoing vertices *u* ∈ *v*_*out*_.

#### Algorithm 3

NextWavefront to compute wavefronts of score s

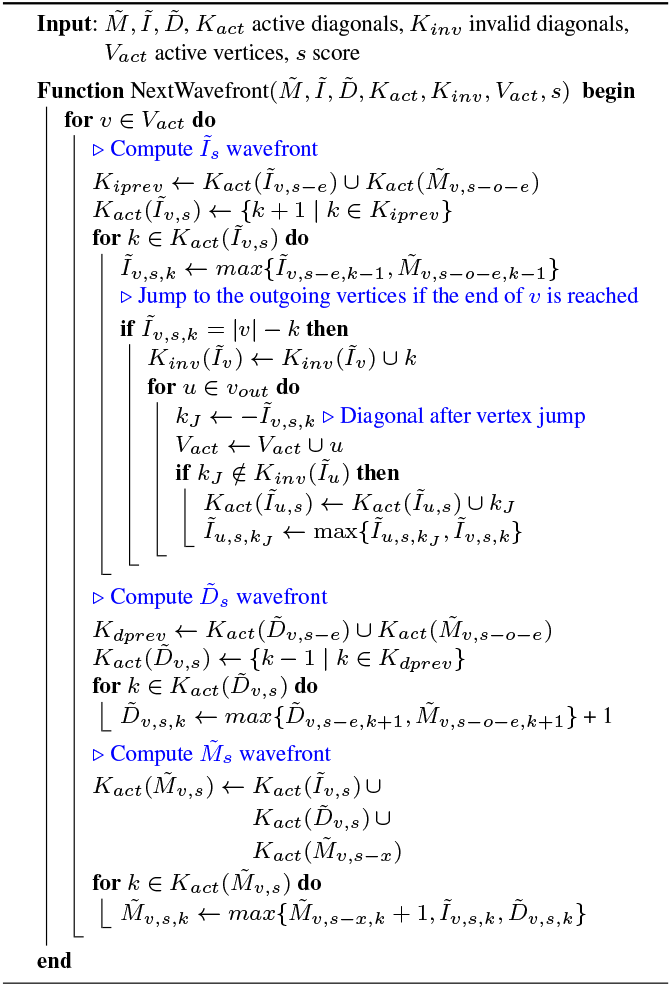

### 3.3 Diagonal Invalidation Strategy

According to Eq. 3, the values on any vertex *v* may depend on wavefront values from adjacent vertices in the sequence-graph. Once a diagonal *k* reaches the end of its associated vertex (i.e., *Ĩ*_*v,s,k*_ = |*v*|−*k* or 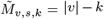, for some score *s*) its offsets propagate to all the outgoing vertices *u* ∈ *v*_*out*_. Thus, due to the score-ascending search of the solution space performed by Theseus, any later exploration of diagonal *k* in vertex *v* will be of a higher or equal score *s*, never leading to better scoring paths. Therefore, Theseus can safely invalidate diagonal *k* in future wavefronts at vertex *v*, avoiding redundant work to compute suboptimal alignments. More in detail, a diagonal *k* can reach the end of a vertex *v* in two distinct situations, noted as *Jump* in Eq. 3. In either of these two cases, Theseus sets the corresponding diagonal as invalid, either for 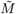 or *Ĩ*. In addition, due to gap propagation on the outgoing vertices, Theseus implicitly invalidates neighboring diagonals every *o* + *e* or *e* scores. This is because the newly invalidated diagonals cannot produce better-scoring paths than those stemming from the associated diagonal on the outgoing vertices.

Theseus compactly represents the invalid diagonals as an ordered set of growing and non-overlapping segments of invalid diagonals. In particular, each segment consists of four values: the start and end of the range [*min*_*diag, max*_*diag*] of invalid diagonals, and the remaining scores *rem*_*down* and *rem*_*up* to invalidate adjacent diagonals due to gap propagation. It is important to note that, given a vertex *v*, the sets of invalid diagonals in 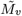, *Ĩ*_*v*_ and 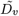 are treated separately, as both *Ĩ*_*v*_ and 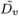 already incorporate the gap opening cost. Theseus initializes segments of invalid diagonals on *jumps*, either on 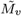 or *Ĩ*_*v*_ . Then, at the end of each iteration on the main loop of Alg. 1, it decreases the scores remaining to expand the segment by one as well. If a remaining value reaches zero, it is set to the default *e* value, and the segment is expanded in the corresponding direction. Then, we sort and merge the updated set to keep it compact and non-overlapping.

This invalidation strategy is applicable to all types of directed graphs, including those with cycles. This is an important feature, as state-of-the-art diagonal invalidation strategies, such as the superbubble pruning implemented in POASTA, are limited to DAGs. Applying those strategies would hinder Theseus’s ability to align sequences when cycles are present in the reference. Overall, our method is designed to operate on general directed graphs, enabling it to accurately model and process repetitive regions in the reference pangenome.

### 3.4 Unified Coordinate System for Efficient Sparse-Wavefront Processing

Due to the divergence of diagonals at outgoing vertices and their convergence at incoming ones, the wavefronts in Theseus are no longer inherently dense but present a sparse behavior. Consequently, we require sparse data structures to represent them, along with novel methods for efficient processing.

Theseus stores alignment information in sparse wavefronts. A sparse wavefront is a set of not necessarily ordered wavefront cells, where each of these cells stores the position of the cell in the corresponding DP matrix and additional information to perform the backtrace of the given alignment problem. However, processing sparse wavefronts introduces two challenges. First, because the cells are non-contiguous, their positions must be tracked individually. Second, the diagonals are often sparsely distributed, potentially unordered, and reused in later sparse wavefront computations, requiring a specialized strategy to process them efficiently. First, to track sparse cells individually, Theseus proposes encoding each cell’s location using its diagonal (*k*), its offset (*off*) within that diagonal, and the vertex (*v*) where it resides.

Second, to efficiently combine sparse wavefronts across vertices and diagonals, we propose using a Unified Coordinate System (UCS). Recall *m* = |*Q*|, the length of the input sequence, and let *n* = max_*v*_ *{*|*v*|*}* be the maximum sequence length within the input sequence-graph *G*. According to Equation 3, the range of possible diagonals that Theseus can potentially explore is [−*m*, · · ·, *n*]. Thus, we introduce an intermediate data structure that maps sparse wavefront data into a common coordinate system of size *n* + *m* + 1, and then converts the result back to a sparse representation.

Figure 2 shows an example of the UCS operation. First ❶, during the *Densification* step, each element of the preceding sparse wavefronts is mapped to its target position in the UCS according to Equation 3, implementing the NextWavefront operation. In this process, the recurrence is applied element-wise, so that each contribution updates its target diagonal. When multiple values map to the same UCS position, we resolve the collision by taking the maximum, as defined by the recurrence 3. Second ❷, *Sparsification* step, once all contributions have been inserted into the UCS, we store the updated diagonals to form the resulting sparse wavefront. Third 3, *Extension* step, the ExtendDiagonal operation is executed on the resulting sparse wavefront. Finally, the UCS is prepared for a new iteration by resetting the modified diagonals to their default values. Figure 2 illustrates this 3-step process.

**Fig. 2:**
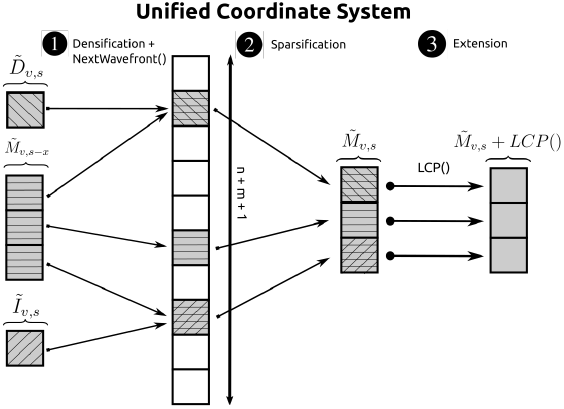
Example of how the Unified Coordinate System (UCS) combines previous sparse wavefronts.

### 3.5 Memory Efficient Sparse-Wavefront Backtrace

Usually, we are interested not only in computing the optimal alignment score but also in recovering the sequence of edit operations that transform *Q* into a sequence spelled by a path in *G*. However, backtracing alignments in a sequence graph requires storing additional information, such as the set and order of vertices visited (i.e., a path in the reference graph).

Backtracing the optimal alignment requires storing the computed results and, once the alignment is finished, reconstructing the set of alignment operations that led to the optimal alignment score. Given its sparse nature, Theseus explicitly stores the position of the incoming cell that leads to the newly created cell. In fact, each cell not only contains its offset value, but also includes data as its vertex number and its diagonal. With all of this information and the starting position of the backtrace, Theseus is able to find the optimal alignment traceback, storing both the traversed path in the reference and the set of edit operations.

However, memory consumption is a key limiting factor for the scalability of any alignment algorithm. For this reason, Theseus takes inspiration from the Singletrack (López-Villellas *et al*., 2025) method to reduce the memory requirements of affine-gap alignment. Singletrack reduces memory consumption by a factor of 3 by observing that the optimal alignment can be recovered from the *M* matrix with minimal overhead during the backtrace stage. Accordingly, algorithms that implement Singletrack avoid storing the *I* and *D* matrices, reducing their memory footprint.

We adapt the Singletrack method to the sequence-to-graph setting in Theseus, extending its memory-efficient backtrace strategy to handle vertex transitions while maintaining optimality. As a result, we reduce memory consumption by storing only the 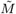 values and the subset of jumping *Ĩ* values. The latter are necessary because long insertions may span multiple vertices and require preserving the corresponding path information.

## 4 Results

We implemented Theseus as a C++ library, publicly available at https://github.com/albertjimenezbl/theseus-lib. All datasets used for experimentation are uploaded in https://zenodo.org/records/18482097.

### 4.1 Experimental Setup

We evaluate the performance of Theseus compared to other state-of-the-art sequence-to-graph libraries and tools. First, in Section 4.2, we evaluate Theseus in the context of computing MSA, comparing it against POA-based libraries such as SPOA (Vaser *et al*., 2017), abPOA (Gao *et al*., 2021), and POASTA (van Dijk *et al*., 2025). Second, in Section 4.3, we evaluate Theseus for pangenome read mapping, comparing its performance against the sequence-to-graph alignment stage of vg map (Garrison *et al*., 2018), along with the alignment kernels of SPOA, abPOA, and POASTA. All experiments are executed single-threaded on an Intel Xeon Platinum 8480 CPU @3GHz node equipped with 256GB of DRAM.

### 4.2 Evaluation on Multiple Sequence Alignment (MSA)

In this section, we evaluate Theseus’s sequence-to-graph alignment algorithm for computing MSA against POA-based libraries such as SPOA, abPOA, and POASTA. It is important to note that all evaluated libraries produce optimal alignments, except abPOA, which uses an adaptive banding strategy to accelerate computations, at the expense of producing suboptimal solutions. For the experiments, each aligner is provided with a set of genomic sequences that are added progressively to the POA graph until all sequences have been processed. We measure the total time and peak memory consumption spent aligning the sequences and updating the corresponding POA graph.

For the evaluation, we follow a methodology similar to that used in the POASTA work. We selected 6 datasets of varying sequence lengths (30Kbp-1Mbp): *SARS-CoV-2* 30Kbp, *M. tuberculosis* 50Kbp, *M. tuberculosis* 250Kbp, *M. tuberculosis* 500Kbp, *M. tuberculosis* 1Mbp and *Mokey pox* 200Kbp. The *SARS-CoV-2* dataset contains 2732 GenBank-complete *SARS-CoV-2* genome assemblies, each approximately ∼30 Kbp in length. The *M. Tuberculosis* 50Kbp, 250Kbp, 500Kbp, and 1Mbp datasets are derived from the same set of 342 RefSeq-complete whole-genome assemblies of *Mycobacterium tuberculosis* genomes by extracting subsequences of different lengths, all starting at the *dnaG* gene. Finally, the *Monkey pox* dataset contains 100 RefSeq-complete whole genome assemblies of *Monkey pox*’s virus, each approximately 200 Kbp in length.

Table 1 shows the execution times of the MSA experiments. We observe that Theseus outperforms the other two optimal aligners, POASTA and SPOA, across all experiments. Specifically, Theseus is between 2.0*×* and 12.0*×* faster than POASTA and between 6.9*×* and 232.2*×* faster than SPOA. Comparing our method to the heuristic abPOA, we observe that while abPOA is 3.0*×* faster on the *M. tuberculosis* 50Kbp dataset, Theseus outperforms abPOA by 6.2*×* on the *SARS-CoV-2* dataset. More importantly, Theseus is guaranteed to always produce optimal alignments, whereas abPOA trades accuracy for speed through heuristic banding and only completes two experiments, running out of memory on datasets with sequences longer than 50 Kbp. In contrast, Theseus is the only method that scales to longer sequences (i.e., 200 Kbp) while maintaining practical runtimes.

**Table 1.** Execution Time (in seconds) of SPOA, abPOA, POASTA, and Theseus on different datasets for the problem of MSA.

| Dataset | SPOA | abPOA | POASTA | Theseus |
| --- | --- | --- | --- | --- |
| SARS-CoV-2 30kbp | 7664 | 205 | 65 | 33 |
| M. tuberculosis 50kbp | 2571 | 124 | 6512 | 370 |
| M. tuberculosis 250kbp | oom | oom | 8690 | 910 |
| M. tuberculosis 500kbp | oom | oom | 29024 | 2453 |
| M. tuberculosis 1Mbp | oom | oom | 71770 | 5996 |
| Monkey pox 200Kbp | oom | oom | timeout | 7785 |

oom: execution exceeded available memory (out of memory).
timeout: execution exceeded 48 hours.

Additionally, Table 2 shows the peak memory consumption of the experiments comparing aligners. The results show that both SPOA and abPOA can only process the two smallest datasets, due to their large memory requirements, requiring 116.9GB and 66.0GB in the *M. tuberculosis* 50Kbp problem, respectively. In contrast, POASTA and Theseus are able to process all datasets and exhibit similar memory usage that scales with the alignment score and problem size, although POASTA exceeds the 48h runtime limit on the *Monkey pox* 200Kbp-long dataset.

**Table 2.**
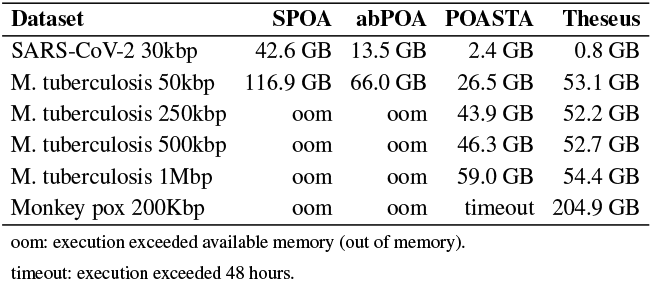
Peak memory consumption (in GB) of SPOA, abPOA, POASTA, and Theseus on different datasets for the problem of MSA.

| Dataset | SPOA | abPOA | POASTA | Theseus |
| --- | --- | --- | --- | --- |
| SARS-CoV-2 30kbp | 42.6 GB | 13.5 GB | 2.4 GB | 0.8 GB |
| M. tuberculosis 50kbp | 116.9 GB | 66.0 GB | 26.5 GB | 53.1 GB |
| M. tuberculosis 250kbp | oom | oom | 43.9 GB | 52.2 GB |
| M. tuberculosis 500kbp | oom | oom | 46.3 GB | 52.7 GB |
| M. tuberculosis 1Mbp | oom | oom | 59.0 GB | 54.4 GB |
| Monkey pox 200Kbp | oom | oom | timeout | 204.9 GB |

oom: execution exceeded available memory (out of memory).
timeout: execution exceeded 48 hours.

### 4.3 Evaluation on Read Mapping

In this section, we evaluate Theseus’ performance for pangenome read-mapping. Pangenome read-mappers, such as vg map from the vg toolkit (Garrison *et al*., 2018), align sequencing reads to a variation graph by combining seeding against a reference graph and sequence-to-graph alignment. The mapping process in vg map starts by finding partial exact matches between the query and the graph, in the form of Maximal Exact Matches (MEMs). Then, these seeds are clustered to identify candidate alignment regions or subgraphs. Finally, vg performs affine-gap sequence-to-graph alignment of the query against those candidate subgraphs. Our evaluation focuses on the sequence-to-graph alignment stage, isolating it from other components of the pangenome read-mapping workflow to enable a fair and informative comparison.

To perform this analysis, we use vg map to extract a collection of sequence-to-graph alignment inputs, consisting of pangenome subgraphs induced from mapping different input sequences to a human pangenome reference (built on top of the GRCh37 reference genome and containing all variation from the Phase 3 of the 1000 Genomes Project (Consortium *et al*., 2015) with an alelle frequency *AF* ≥ 1%). For each input read mapped to the forward strand, we use vg map to identify clusters of Maximal Exact Matches (MEMs) and induce a candidate subgraph. As a result, we obtain a collection of sequence-to-graph alignment instances, each defined by an induced subgraph, an alignment sequence, and the starting mapping position in the graph. We modify the induced subgraph by adding an additional node pointing to the starting position to enable a fair comparison between the different methods. Then, we evaluate the performance of different aligners to compute the alignment of the input sequence to the induced subgraph.

We evaluate the alignment kernels of vg map (Garrison *et al*., 2018), SPOA, abPOA, POASTA, and Theseus on three short read datasets of lengths 100bp, 150bp, and 250bp. The Illumina 100bp dataset comes from individual HG00096 of the 1000 Genomes Project and the other two were extracted from the Genome in a Bottle project (Zook *et al*., 2016). We measure alignment throughput by each sequence-to-graph aligner across the selected datasets.

Figure 3 compares the alignment throughput of vg map, SPOA, abPOA, POASTA, and Theseus. We observe that Theseus is between 1.9*×* and 16.9*×* faster than the other alignment kernels. Note that the performance of Theseus is sensitive to the alignment difficulty (i.e., alignment score) of the input sequences across the different datasets and sequencing technologies. This experiment serves as a proof of concept to assess Theseus in a realistic read-mapping setting, without aiming to replace the full pangenome mapping pipeline.

**Fig. 3:**
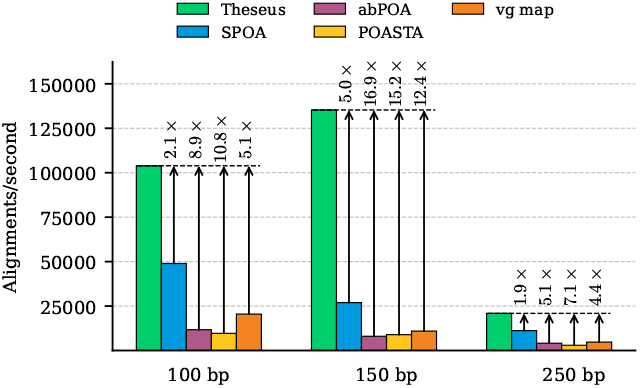
Alignment throughput of Theseus compared to other aligners on three datasets of 100bp, 150bp, and 250bp.

## 5 Discussion

In this work, we present Theseus, a fast, scalable, and affine-gap sequence-to-graph aligner. Theseus exploits the expected high similarity between genomic sequences to accelerate alignment and reduce its memory footprint. Compared to other alternatives, Theseus naturally supports cycles, making it adequate to handle all sorts of directed graphs.

We introduce several algorithmic innovations to cope with the requirements of this new framework. First, we extend the affine-gap recurrence equations from the original algorithm to work with graph-like sequence references. Second, we implement a diagonal invalidation strategy that tracks previously processed and implicitly explored diagonals to avoid redundant computation. Third, we introduce the notions of sparse wavefronts and a Unified Coordinate System to handle the non-contiguous data arising from graph alignment. Finally, we introduce a memory-efficient backtrace algorithm, based on the recently published Singletrack (López-Villellas *et al*., 2025) approach to backtrace in pairwise alignment, tailored to the sparsity of sequence-to-graph alignment.

We evaluate Theseus on two distinct use cases, highlighting its potential as a general sequence-to-graph alignment tool. In the Multiple Sequence Alignment (MSA) problem, we observe that Theseus consistently outperforms both POASTA (van Dijk *et al*., 2025) and SPOA (Vaser *et al*., 2017) across all tested datasets, showing a 2.0*×* to 232.2*×* speed-up. We also compare it with abPOA, a heuristic POA aligner that accelerates computation at the expense of optimality. We demonstrate that Theseus is 3.3*×* faster on average, while guaranteeing optimality. We also observe that Theseus is the only tool that scales to larger problems while maintaining manageable runtimes. In the Read Mapping use case, intended as a proof of concept of Theseus’s capabilities, we compare Theseus with the alignment kernels of vg map, SPOA, POASTA, and abPOA on short reads. We observe that Theseus is between 1.9*×* and 16.9*×* faster than the other aligners.

Overall, Theseus bridges the gap between optimality and scalability in sequence-to-graph alignment. By combining output-sensitive exploration with sparse wavefront processing, it enables affine-gap alignment on large and cyclic graphs at a practical cost. As a result, we expect Theseus to provide a suitable building block for next-generation MSA pipelines and scalable pangenome analysis as reference graphs and datasets continue to grow in the future to come.

